# State-specific Peptide Design Targeting G Protein-coupled Receptors

**DOI:** 10.1101/2024.11.27.625792

**Authors:** Yang Xue, Hong Wang, Jun Li, Jianguo Hu, Zhiyuan Chen, Zhi Zheng, Lihang Liu, Kunrui Zhu, Jingzhou He, Huanzhang Gong, Xiangqun Li, Xiaonan Zhang, Xiaomin Fang

## Abstract

G protein-coupled receptors (GPCRs) play critical roles in various physiological processes and serve as pivotal targets for drug discovery. Peptides are particularly compelling therapeutic agents for targeting GPCRs, as they frequently exhibit superior affinity, selectivity, and potency compared to small-molecule drugs. However, the states (active and inactive) of GPCRs profoundly influence their interactions with ligands, highlighting the need for state-specific strategies in peptide design. To address this, we developed an efficient state-specific peptide design pipeline targeting GPCRs. This framework enables the tailored design of agonists or antagonists based on the active or inactive conformational states of GPCR structures. A state-specific folding model, optimized for GPCR-peptide complexes, was used to select high-potential peptides from the pool of designed candidates. Using this approach, we successfully identified both agonist and antagonist peptides for the Apelin Receptor (APJR) and the Growth Hormone Secretagogue Receptor (GHSR), as well as a competitive inhibitor for the Glucagon-Like Peptide-1 Receptor (GLP-1R).

## Introduction

G-protein-coupled receptors (GPCRs) are a large and diverse group of membrane receptors that play crucial roles in cellular signal transduction. Their involvement in various physiological processes makes them attractive targets for therapeutic intervention ^1^. Despite the success of small molecules and biologics in modulating GPCR activity, these approaches often face limitations such as off-target effects ^2^. Peptide-based therapeutics offer a promising alternative due to their remarkable pharmacodynamic properties: high affinity, selectivity and potency ^3;4^, though their clinical use is challenged by limited in vivo stability and complex synthetic requirements. These limitations can be mitigated through strategies such as incorporating non-natural amino acids or chemical modifications ^5^ to enhance stability and synthetic feasibility.

Conventional methods for developing peptide drugs involve leveraging natural peptides and hormones found in the human body, as well as creating peptides that mimic these hormones ^6–8^. Peptides can also be sourced from natural products, including those derived from bacteria, fungi, and plants ^9–13^. Another approach involves the use of phage display techniques to discover and optimize peptide candidates ^14^.

Recently, AI-based peptide rational design has gathered increasing attention ^15–17^. EvoBind ^18;19^ employs in silico-directed evolution to design peptides and then evaluates the designs using confidence scores from the folding model AlphaFold-Multimer ^20^. EvoPlay ^21^ improves it by replacing directed evolution with Monte Carlo tree search (MCTS) and reinforcement learning policy ^22^, leading to more data-efficient peptide design. Additionally, PepMLM ^23^ uses a protein language model (PLM) to design peptide binders. Approaches for designing mini-proteins may also be applicable to peptide designs, such as AlphaFold-2 hallucination ^24^, RFDiffusion-MPNN ^24^, AlphaProteo ^25^ and BindCraft ^26^.

**Figure 1.**
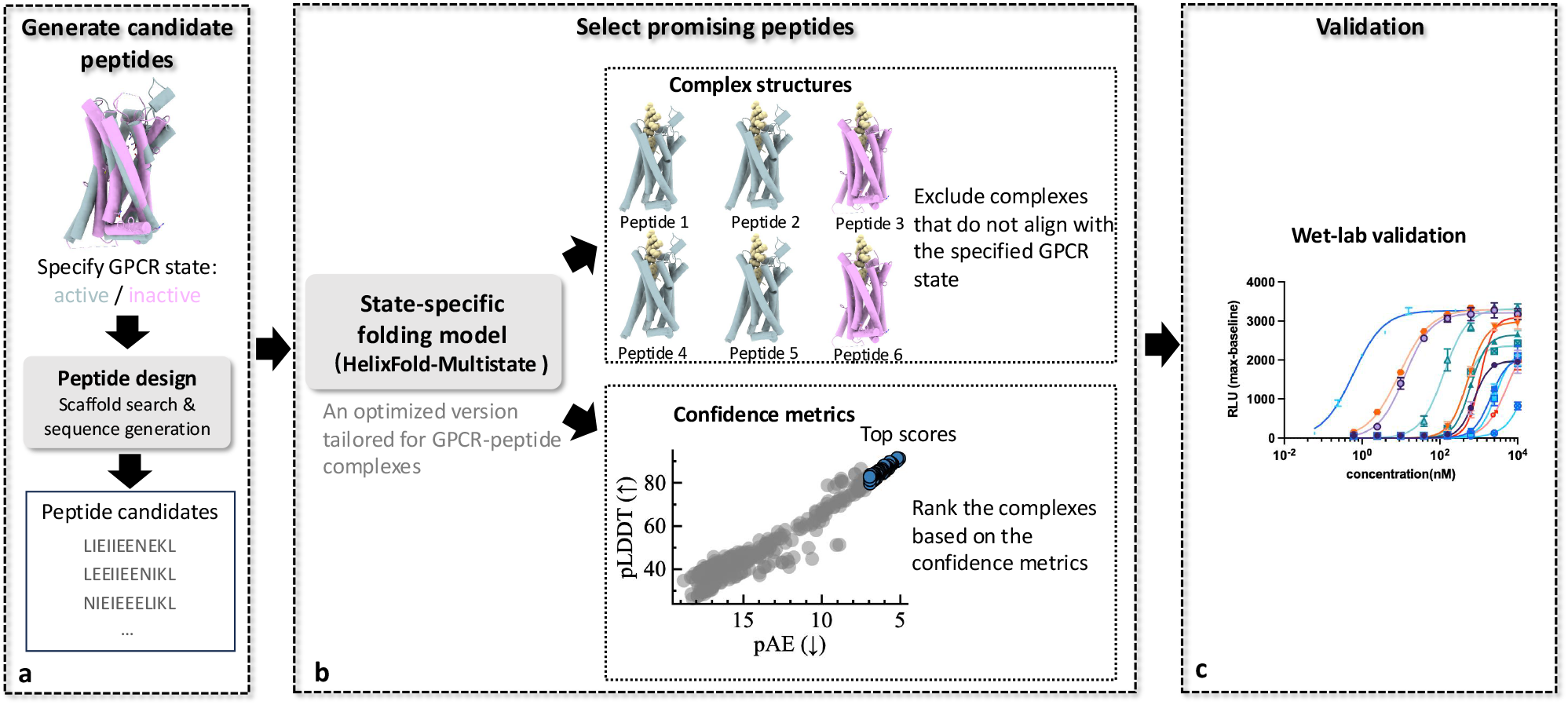
State-specific peptide design pipeline. **(a)** Generation of the candidate peptides through the peptide design modules. **(b)** State-specific folding model (HelixFold-Multistate), an optimized version tailored for GPCR-peptide complexes, to select promising peptides. **(c)** Wet-lab experimental validation.

Although AI-driven peptide design approaches have successfully generated high-affinity binders, targeting GPCRs presents unique challenges due to their dynamic conformational landscape and functional plasticity as shown in GPCRmd^27^. GPCRs can adopt multiple active and inactive states, each stabilized by distinct ligand interactions, which determine whether a peptide functions as an agonist or antagonist. Effective GPCR-targeting peptides must not only achieve high binding affinity but also selectively stabilize specific receptor conformations, thereby modulating the energy landscape to favor desired functional outcomes. Therefore, integrating GPCR conformational states and state-specific interactions into peptide design strategies is critical for achieving precise modulation of receptor activity. Existing approaches for peptide design targeting GPCRs, such as HelixDiff ^28^, typically define the structure of the GPCR protein used in the generation model. However, these methods do not allow for precise control over the GPCR’s functional state (e.g., active or inactive). This limitation hinders the ability to design peptide agonists and antagonists that accurately correspond to specific receptor states.

In this study, we present a computational method for state-specific peptide design targeting GPCRs (Figure 7). The workflow begins with the generation of a diverse pool of candidate peptides using a peptide design module ^29;30^ (Figure 7a). A state-specified folding model (Figure 7b) is then employed to define the active or inactive states of GPCR targets, predict the structures of GPCR-peptide complexes, and assign a confidence score to each complex. Complexes that do not align with the specified GPCR state, such as those with the binding pocket located within the membrane or those exhibiting a significant RMSD in the state-specific TM domain compared to the expected state, are filtered out. The remaining complexes are then ranked based on the integrated application of multiple confidence metrics. This ranking process allows for the prioritization of peptide candidates with the highest likelihood of success for further validation through in silico and experimental methods (Figure 7c). To refine the folding model for GPCR-peptide interactions, we fine-tuned HelixFold-Multimer (HF-Multimer) ^31^, a reproduction of AlphaFold-Multimer (AF-Multimer) ^20^ on GPCR-peptide ^32^ and protein-peptide ^33^ datasets. This refinement resulted in improved predictive accuracy for GPCR-peptide complexes. Furthermore, we incorporated state-specific modeling techniques inspired by AlphaFold-Multistate (AF-Multistate) ^34^, enabling precise differentiation between the active and inactive states of GPCRs. These advancements facilitate the rational design of peptide agonists and antagonists specifically tailored to target specific GPCR receptors. The resulting model, named HelixFold-Multistate (HF-Multistate), provides a robust computational tool for advancing the design of state-specific peptide therapeutics targeting GPCRs.

We validated that the GPCR-peptide structures predicted by HelixFold-Multistate are more effective at preserving the correct activation states of GPCRs while providing more accurate GPCR-peptide interaction poses for both active and inactive states of GPCR targets. Furthermore, the structural confidence scores derived from HelixFold-Multistate demonstrated a strong correlation with experimental affinity data for both peptide agonists and antagonists. We tested the state-specific peptide design pipeline by designing both agonists and antagonists for three GPCR targets, Apelin Receptor (APJR) and Glucagon-Like Peptide-1 Receptor (GLP-1R), and Growth Hormone Secretagogue Receptor (GHSR) using in-silico and experimental validation. Designing antagonistic peptides for GPCRs is likely more challenging than agonists as most existing peptide therapeutics for GPCRs are agonists ^4^. Additionally, while the GPCRdb database ^32^ contains few antagonist peptides, most GPCR-targeting antagonists are small molecules. The pipeline successfully generated de novo high-affinity agonists (<100 nM) and moderate-affinity antagonists, highlighting the precision and flexibility of the proposed methodology. Notably, the agonist peptides targeting APJR exhibit a high affinity, with an EC50 < 10 nM. The antagonist peptides targeting GLP-1R have an IC50 of 874 nM.

## Results

We first evaluated the structural prediction accuracy and screening effectiveness of the optimized state-specific folding model, HF-Multistate, which serves as the most critical component of our peptide design pipeline. Following this, we assessed the pipeline’s performance by designing peptides for three GPCR targets, aiming to evaluate its ability to generate target-specific agonists and antagonists.

### GPCR-Peptide Structure Prediction Capacity

We evaluated the performance of different folding models from two perspectives: the accuracy of GPCR-peptide interface predictions and the structural accuracy of GPCR states, particularly in key regions. Our optimized state-specific folding model, HelixFold-Multistate (HF-Multistate), was benchmarked against AlphaFold-Multimer (AF-Multimer) ^20^, HelixFold-Multimer (HF-Multimer) ^31^ and AlphaFold-Multistate (AF-Multistate) ^20^. It is worth noting that AF-Multimer and AF-Multistate utilize an ensemble of five models, each executed five times with different random seeds for each prediction. In contrast, HF-Multimer and HF-Multistate achieve competitive results using a single model executed five times with varying seeds.

We curated a recent dataset of agonist and antagonist peptide-GPCR structures from GPCRdb^32^, applying cutoffs for both release date and GPCR type. This process guarantees no overlap in GPCR types between the evaluation and training sets. Further details on the evaluation set are available in Supplementary Section **A**. We employed multiple evaluation metrics, including DockQ ^35^, interface root-mean-square deviation (iRMS, a component of DockQ), % Correct (DockQ > 0.8), % Correct (DockQ > 0.49), and % Correct (iRMS < 2.0), to comprehensively assess the performance of various folding models in predicting GPCR-peptide interfaces. DockQ is a composite metric ranging from 0 to 1 that evaluates the quality of predicted protein–protein or protein–peptide interfaces by combining interface RMSD (iRMSD), ligand RMSD (lRMSD), and the fraction of native contacts (Fnat). Higher scores indicate better interface accuracy. iRMS (interface RMSD) measures the deviation between the C! atoms of interface residues in the predicted and experimental structures, focusing specifically on the local binding interface. All evaluation metrics were computed with respect to the experimentally resolved GPCR–peptide complex structures. The model was conditioned on the specified receptor state, which was derived from the annotations in the experimental data. As shown in Figure 8(a-c), HF-Multistate demonstrates top-tier performance in predicting GPCR–peptide interactions, showing a noticeable advantage over AF-Multistate. While AF-Multistate has been demonstrated to accurately predict the activation states of monomeric GPCRs^34^, it faces limitations in predicting interactions within GPCR-peptide complexes. The comparison between HF-Multistate and AF-Multistate highlights the importance of incorporating GPCR-peptide complex structural data for fine-tuning, as such data can improve predictive accuracy, particularly in modeling state-specific interactions.

Although DockQ and related metrics are well suited for evaluating the accuracy of peptide–GPCR interfaces, they are less sensitive to structural precision in functionally important regions of the GPCR receptor, i.e., the transmembrane (TM) domain ^34^. To address this limitation, we conducted a focused analysis of TM domain structural accuracy across two major GPCR families: Class A (rhodopsin-like receptors) and Class B (secretin-like receptors). Following the methodology outlined in a previous study ^36^, we focused our analysis on TM3 and TM6 for Class A GPCRs, on TM3, TM6, and TM7 for Class B GPCRs, since these regions are crucial for peptide hormone recognition and receptor activation. In Class A GPCRs, TM3 and TM6 undergo state-dependent conformational changes: TM6 moves outward in active states to facilitate G-protein binding. At the same time, TM3 exhibits a cytosolic tilt and overall rotation with only minor changes. In Class B GPCRs, TM3, TM6, and TM7 exhibit more intricate state-specific dynamics: TM6 undergoes outward shifts to drive receptor activation, TM3 maintains structural integrity through minor adjustments, and TM7’s extracelluar region undergoes significant conformational changes to enable peptide hormone binding and activation. Across both classes, TM3 and TM6 function as key ‘macroswitches’ in the receptor activation process. Figures 8(d-e) and (f-h) present a comparative analysis of the performance of multiple folding models on TM domains for Class A and Class B GPCRs, respectively, based on the root-mean-square deviation (RMSD) of key transmembrane (TM) domains. RMSD quantifies the structural deviation between predicted and experimental atomic coordinates, providing a measure of accuracy in modeling these functionally important regions. For Class B GPCRs, the comparison of % Correct (TM RMSD < 2.0) is not very significant. Therefore, we included the distribution of all data points for clarity. AF-Multistate and HF-Multistate outperform AF-Multimer in structural prediction accuracy across the TM domains, with notable improvements in the critical TM3 and TM6, emphasizing the advantages of state-specific modeling. More importantly, HF-Multistate consistently achieves the highest precision across all key TM domains in terms of TM RMSD, highlighting HF-Multistate’s superior ability to incorporate the distinct activation states of GPCRs during peptide binding, providing a deeper understanding of their functional dynamics.

**Figure 2.**
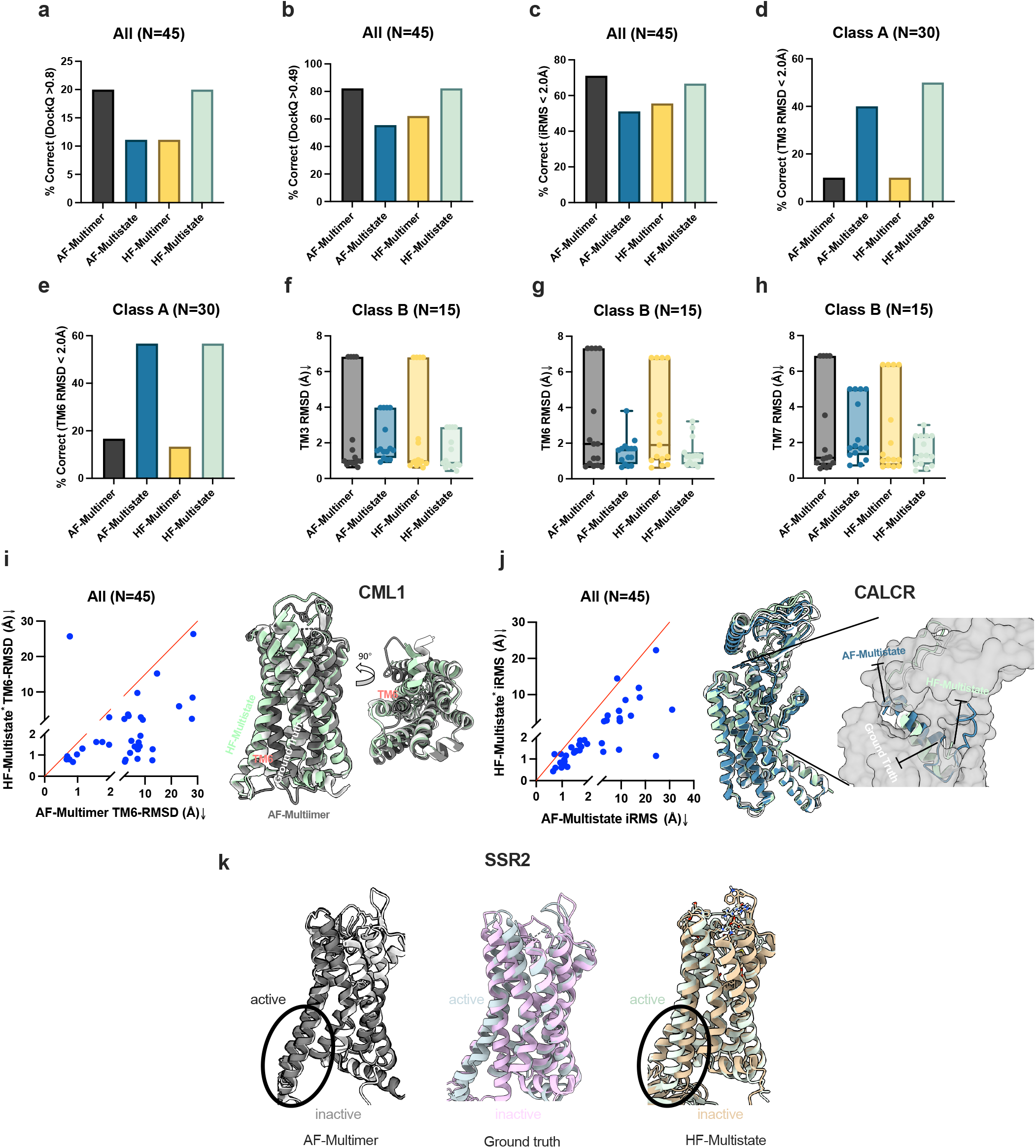
Comparison of folding models for structural prediction accuracy in recent GPCR-peptide complexes from the PDB. **(a-c)** Comprehensive assessment of GPCR-peptide interface prediction accuracy using multiple evaluation metrics. **(d-e)** Assessment of structural accuracy in GPCR States through evaluation of the transmembrane (TM) domain in GPCR class A. **(f-h)** Assessment of the TM domain in GPCR class B. **(i)** Scatter plot comparison of TM6-RMSD between AF-Multimer and HF-Multimer, indicating HF-Multistate is more state-sensitive than AF-Multimer. CML1 active mode (PDB ID: **7YKD**) is taken as an example to illustrate that AF-Multimer (Gray) predicts wrong activation state while HF-Multistate (Light Green) can predict correct state. d"id,8F0K**8F0K) is taken as an example.(j)** Scatter plot comparison of iRMS between AF-Multistate and HF-Multistate for GPCR Class B. CALCR active mode (PDB ID: **8F0K**) is taken as an example. d"id,7XAU**7XAU) and inactive mode (7XNA) while HF-Multistate (Right) can.(k)** Illustrate state-specific feature of HF-Multistate using SSR2, AF-Multimer model (Left) cannot distinguish receptor state from active mode (**7XAU**) and inactive mode (**7XNA**) while HF-Multistate (Right) can.

To demonstrate HF-Multistate’s improved modeling of functionally relevant GPCR regions over both AF-Multimer, we selected cases based on TM6 RMSD (Fig. 8i). In the resulting scatter plots, each point represents a complex from the test set, with the x-axis and y-axis showing the same metric calculated by HF-Multistate and AF-Multimer, respectively. Similarly, to emphasize HF-Multistate’s improvements in interface accuracy over AF-Multistate, we performed a one-to-one comparison of these two methods using iRMS (Fig. 8j). To provide a more detailed comparison, we analyzed three representative cases: CML1, CALCR and SSR2. For CML1 in Figure 8i, HF-Multistate accurately predicts the CML1 activation state, especially the TM6, whereas AF-Multimer erroneously identifies an incorrect GPCR state. For CALCR in Figure 8j, while AF-Multistate identifies the correct binding pocket, it misorients the peptide, causing the N-terminus to face upward, a mistake not observed with HF-Multistate. For SSR2 in Figure 8k, the agonist peptide **FCFWKTCT** and the antagonist peptide **FCYWKTCY** are challenging to differentiate due to their subtle sequence differences. AF-Multimer (Left) fails to distinguish between the active and inactive complex states, whereas HF-Multistate (Right) successfully differentiates the two states. These findings highlight the inherent challenges in predicting GPCR-peptide complex structures and underscore the importance of refining models with relevant structural data to achieve improved predictive accuracy.

### Ability to Select Promising Peptides Targeting GPCRs

Identifying peptide candidates with both strong binding affinity and structural stability is crucial for the effective design of GPCR-targeted peptides. To evaluate whether HF-Multistate’s confidence scores can aid in this selection process, we assessed their correlation with experimentally measured affinity data from the PPI-Affinity dataset focused on peptides binding the CXCR4 receptor ^37^.

We first analyzed the relationship between the peptide average pLDDT (predicted Local Distance Difference Test) and the GPCR-peptide average pAE (predicted Aligned Error), as indicated by AF-Multistate and HF-Multistate, in comparison with the binding affinity. The pLDDT, introduced by AlphaFold2^38^, is a per-residue confidence metric ranging from 0 to 100, where higher values indicate greater structural accuracy and stability in predicted peptide conformations. Similarly, pAE, as described in AlphaFold-Multimer ^20^, quantifies the expected error in residue-residue alignments, with lower values reflecting higher confidence in binding interface predictions. According to their physical interpretation, a higher pLDDT suggests a more structurally plausible, potentially more natural and stable peptide sequence, while a lower pAE indicates greater confidence in binding predictions due to more reliable alignment distances. Although pLDDT and pAE were initially designed to assess structural confidence, our analysis reveals a weak correlation between these metrics and GPCR-peptide binding affinity (Figure 9a and Figure 9b). Notably, the correlation between HF-Multistate’s predicted metrics and binding affinity is more potent than that of AF-Multistate, suggesting that HF-Multistate provides a more informative structural assessment in this context. This improved correlation indicates its potential to enhance peptide ranking for experimental validation.

To further assess the predictive utility of the multistate models, we examined whether their confidence metrics—pAE and pLDDT—can differentiate peptide functionality across GPCR activation states. The CXCR4–peptide affinity dataset used in this study focuses on inhibitory peptides, where the inactive receptor state represents the correct functional conformation. For each peptide–receptor pair, we compare confidence scores of the correct (inactive) and incorrect (active) receptor states. Samples were classified into three groups: Correct-favored (significantly higher confidence for the correct state), Neutral (similar confidence scores for both states), and Incorrect-favored (higher confidence for the incorrect state). Our results show that HF-Multistate produces a notably higher proportion of Correct-favored predictions, indicating it more reliably distinguishes peptide functionality by favoring the biologically relevant receptor conformation. Conversely, AF-Multistate predictions cluster mainly in the Incorrect-favored group for pAE and Neutral group for pLDDT, demonstrating limited sensitivity to receptor state and reduced ability to capture functional differences (Figure 9c and Figure 9d).

**Figure 3.**
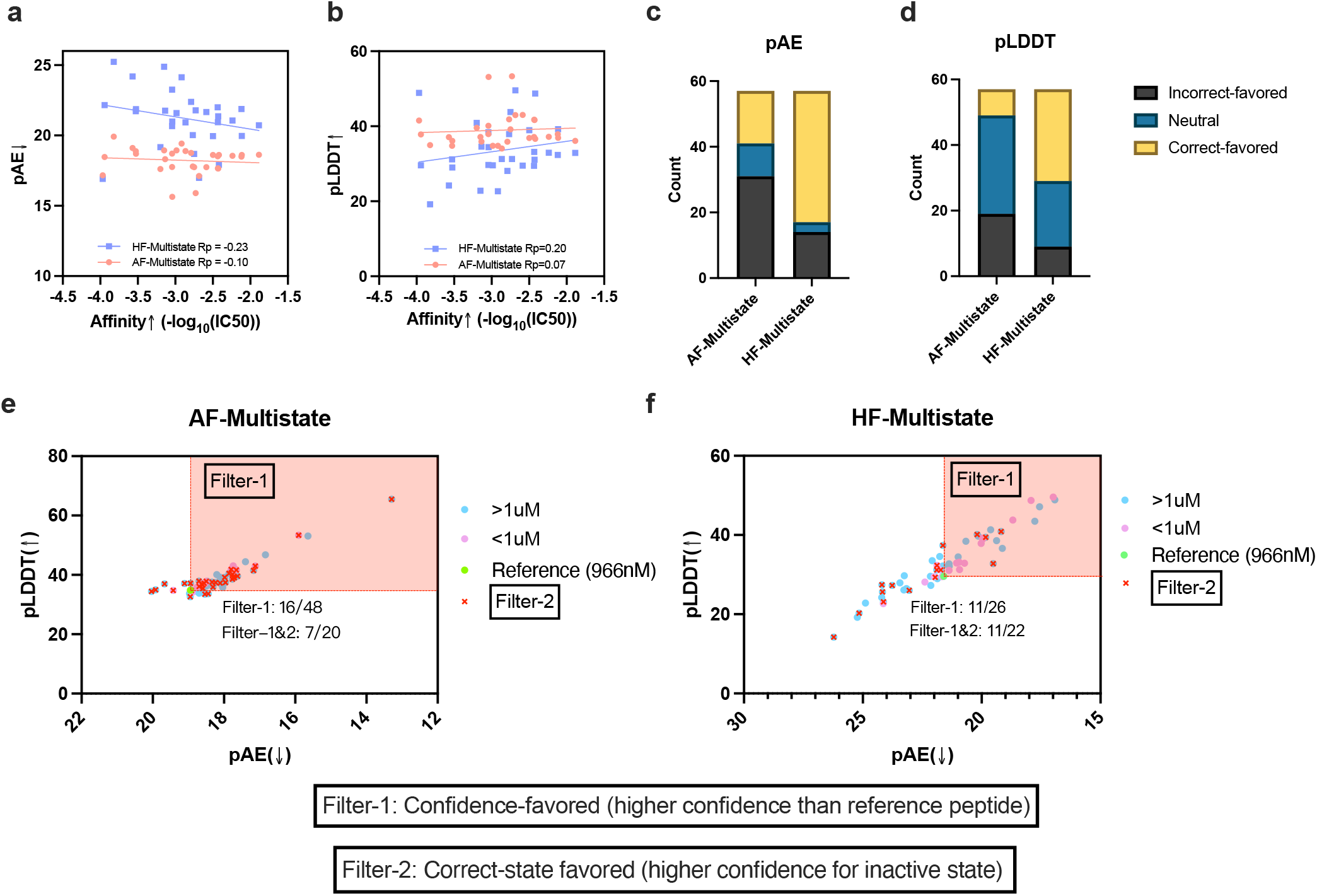
Comparing the screening ability of metrics outputted by AF-Multistate and HF-Multistate on CXCR4-peptide affinity data. (**a, b**) Pearson correlation analysis of pAE and pLDDT scores with peptide affinity, respectively. The correlation between the metrics of HF-Multistate and experimentally determined affinities is higher than that of AF-Multistate. (**c, d**) Confidence-based distinction between GPCR activation states. HF-Multistate shows a stronger tendency to favor the correct state (inactive) with higher confidence, as reflected in both pAE and pLDDT metrics. In contrast, AF-Multistate often fails to differentiate between states, or assigns higher confidence to the incorrect state (active). **(e, f)** Filter rules for screening peptides with a possible higher affinity than the reference peptide with an IC50 of 966nM. The region that satisfies Filter-1 is highlighted in red (Filter-1, containing peptides with confidence scores higher than the reference peptide). Peptides that do not satisfy Filter-2 (the confidence score for the correct state is expected to be higher than that for the incorrect state) are marked with a red cross marker, while peptides that satisfy Filter-2 are represented with other markers.

The preceding analysis (Figure 9(a-d)) indicates that HF-Multistate’s confidence metrics can identify peptides with high affinity and functional relevance for GPCR targeting from multiple perspectives. To further evaluate the combined effectiveness of these predictive capabilities, we applied two filtering rules: Filter 1: Confidence-favored — candidates must have both pLDDT and pAE scores superior to those of the reference peptide, indicating higher overall prediction confidence. The reference peptide is from the CXCR4-peptide affinity dataset with a reported IC50 of 966nM, and was selected as a reference because its affinity is close to the 1µM threshold used to define hits. Filter 2: Correct-state favored — candidates must show stronger model confidence when conditioned on the correct receptor state (inactive), i.e., pAE_inactive_ < pAE_active_ or pLDDT_inactive_ > pLDDT_active_. A comparison of AF-Multistate and HF-Multistate (Figure 9(e-f)) shows that after applying Filter 1, HF-Multistate achieves a hit rate of 11/26 (42%), which is significantly higher than AF-Multistate’s 16/48 (33%). After applying both Filter 1 and Filter 2, the hit rate for HF-Multistate increases to 11/22 (50%), while AF-Multistate’s hit rate is 7/20 (35%). Notably, AF-Multistate performs only slightly better than the random predictor, which has a hit rate of 17/56 (30%). Here, a hit is defined as a peptide with an affinity below 1 µM. These results demonstrate the enhanced capability of HF-Multistate in identifying functionally relevant peptides through a multi-criteria filtering approach.

### Validation on APJR

**Figure 4.**
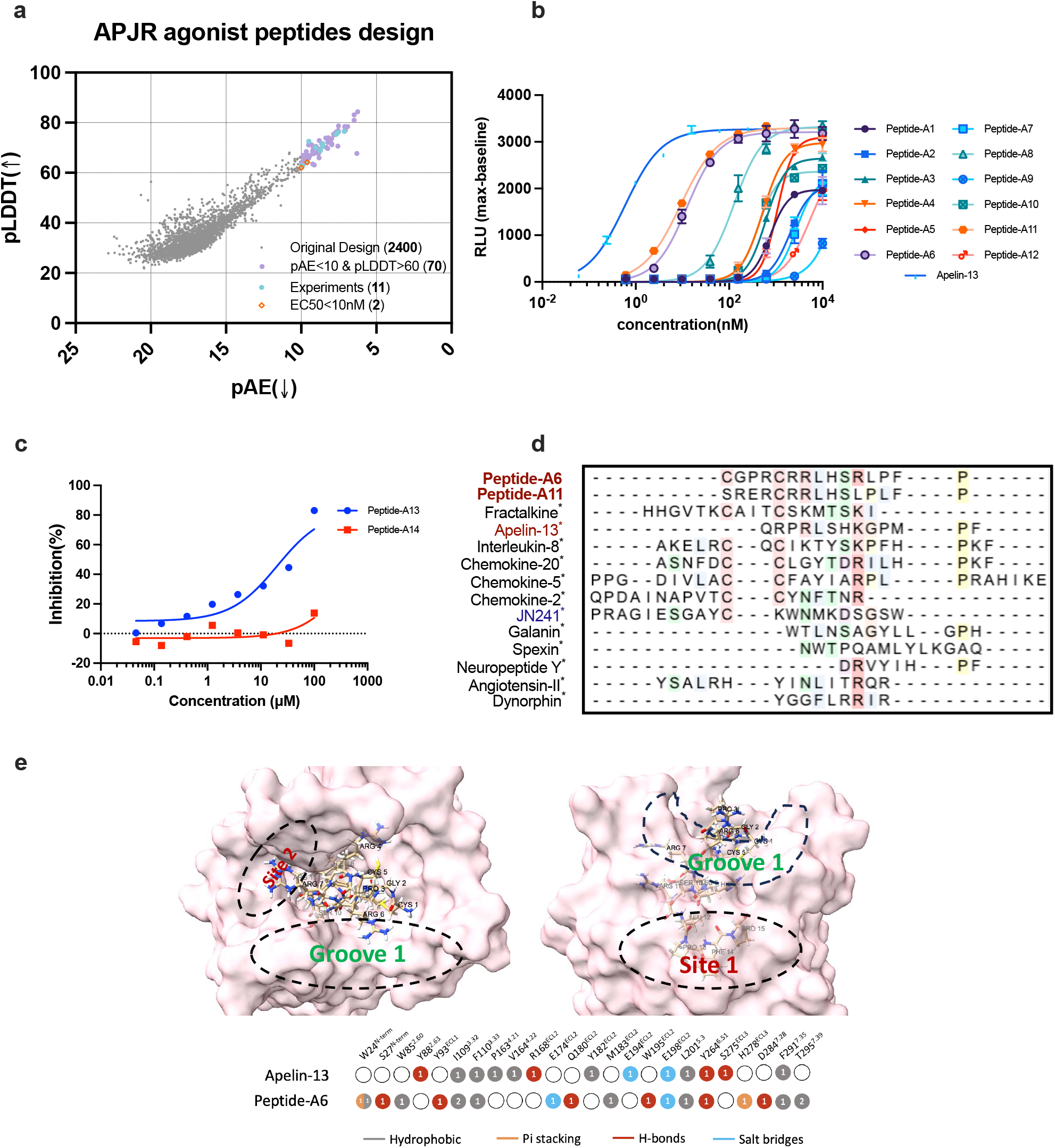
Peptide filtering and analysis for APJR. **(a)** Multiple stages of computational screening. The distribution of pAE and pLDDT scores for the designed peptides is shown across various stages. **(b)** Dose-response curves for the tested peptides, showing relative luminescence units (RLU) as a function of peptide concentration (nM). **(c)** Inhibition ratios (%) of APJR activity in response to increasing concentrations of designed APJR antagonist peptides. **(d)** Sequence alignment of Peptide-A6 and Peptide-A11 with top seeds of the peptide backbone (marked with ^*^) used in the design pipeline. Agonist peptides targeting APJR are indicated in red, while antagonist seeds are shown in blue. **(e)** Top and lateral views of Peptide-A6 and APJR complex predicted via HelixFold-Multistate. Bottom part is comparison of interaction patterns of Apelin-13 (PDB ID: **8XZG**) and our design Peptide-A6.

We begin by evaluating our design pipeline using the Apelin Receptor (APJR), a multifunctional G protein-coupled receptor (GPCR) involved in cardiovascular regulation, fluid homeostasis, metabolism, and angiogenesis ^39^. Our objective is to design both agonists and antagonists that target the APJR.

**Table 1.**
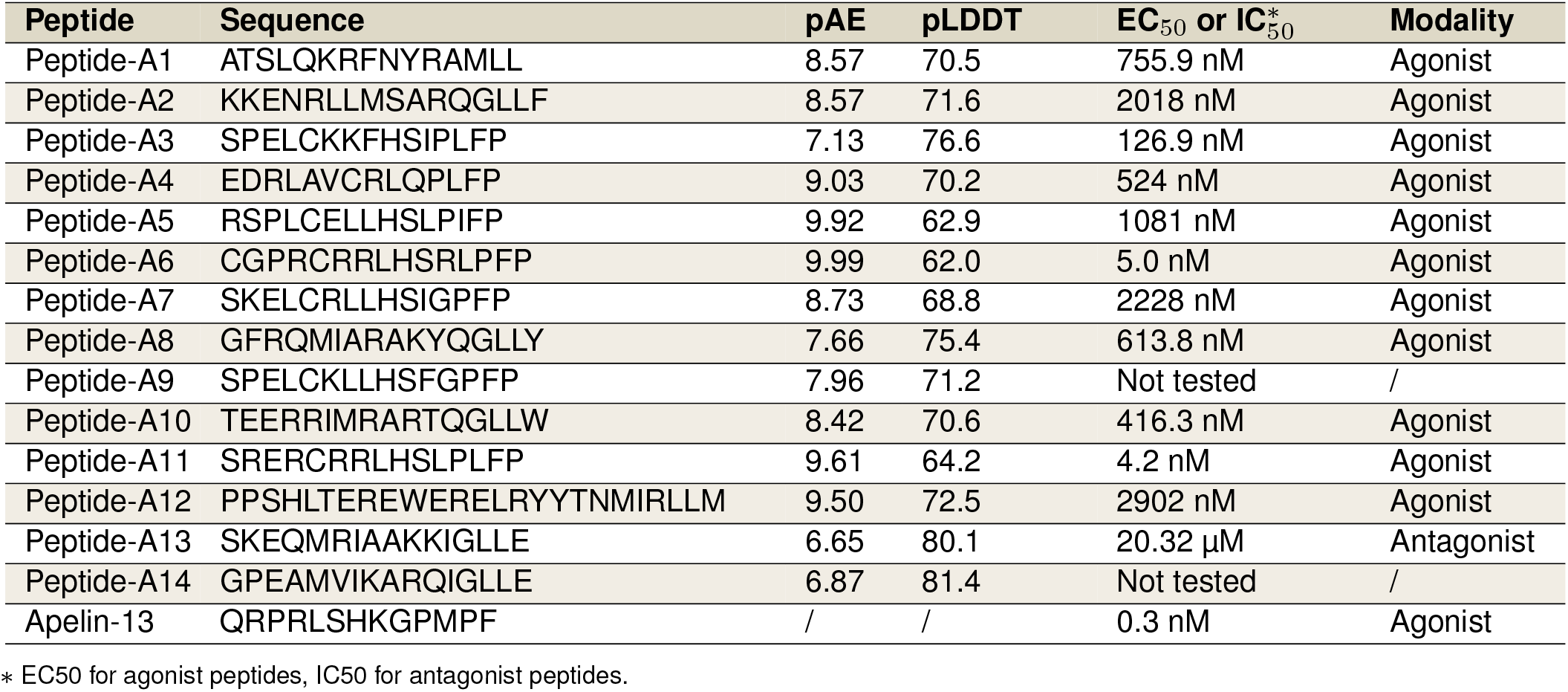
Experimental affinity and peptide modality for designed peptides targeting APJR

For agonist design, we first employ FoldSeek ^40^, a computational tool for rapid and sensitive structural similarity searches, to identify peptide backbones. We chose FoldSeek over manual selection to efficiently and automatically retrieve a structurally diverse set of peptide–GPCR complexes across targets. The Supplementary Section **C** describes the peptide seeds discovery process using FoldSeek for APJR. Table **S1** and **S2** list the peptide seeds used for agonist and antagonist peptides design respectively. We screen for peptides and protein fragments that interact with proteins structurally similar to APJR, serving as candidate scaffolds. Subsequently, a diverse library of 2,400 peptide sequences is generated using a protein inverse folding model ^29^, which produces peptide sequences to fit the identified backbone structures while accommodating the receptor’s binding interface. The designed peptide sequences undergo multiple stages of computational screening (Figure 10a). HF-Multistate is employed to predict the structural conformations of APJR and the peptide candidates, alongside their structural confidence metrics. Based on these evaluations, 70 candidates with pLDDT > 60 and pAE < 10 are selected for further assessment. From this subset, 11 peptides are selected for synthesis and in vitro binding affinity assays. The CHO-K1/AGTRL1/G!15 cell line is used to express the APJ receptor, and a calcium mobilization assay is performed to quantify receptor activation. A similar strategy is applied for antagonist design.

We first assess whether multiple stages of computational screening effectively identify high-quality peptides. Based on Table 4, all tested peptides exhibit activity toward APJR at concentrations below 100 µM, indicating that HF-Multistate’s confidence metrics can effectively identify peptides with potential APJR-binding capability. However, the peptides with the highest affinity, A6 and A11 (highlighted with red diamonds with EC50 values less than 10 nM), do not correspond to the highest pLDDT and pAE scores within the experimental set. This suggests that structural confidence metrics alone are insufficient for accurately ranking binding affinity, highlighting the need to incorporate computational energy-based evaluations, such as MMGBSA, for more precise affinity differentiation. However, simulating the structure of GPCR-peptide complexes with a membrane can be time-consuming. In this study, we employed a coarse simulation without a membrane to screen and select top peptides for subsequent wet-lab experiments.

We next examine the experimental validation of the designed peptides, analyzing their dose-response behavior and modality. The corresponding curves for agonists and antagonists are presented in Figures 10b and 10c, respectively. The Relative Light Unit (RLU) indicates the level of receptor activation and response to the peptides. The inhibition ratio measures the percentage of inhibition of the receptor’s activity by the peptides. In the agonist design, two peptides demonstrated EC50 values below 10nM, whereas the antagonist design yielded a peptide with an IC50 of 20.3µM.

For comparison, Apelin-13^41^, a naturally occurring APJR agonist critical for cardiovascular and metabolic regulation, serves as a reference control for agonist activity (Figure 10b). Although the designed peptides exhibit weaker activity compared to Apelin-13, their sequences differ significantly from Apelin-13 as shown in the sequence alignment results in Figure 10d using MAFFT^1^, suggesting a potentially distinct binding mode. To gain deeper insights into the interaction patterns, we examined the predicted interaction between Peptide-A6 and APJR, as illustrated in Figure 10e. In comparison with previous studies on AMG3054^41^, ELA-32^42^ and Apelin-13^43^, Peptide-A6 is found to interact not only with site 1 (F110^3.33^, Y264^6.51^, W85^2.60^, etc.) and site 2 (W24^N-term^, D284^7.28^, etc.), but also with groove 1 (E174^ECL2^, W195^ECL2^, etc., all in ECL2). Importantly, these residues have been experimentally validated in prior studies, providing additional support for the accuracy of our HF-Multistate predictions in representing the Peptide-A6-APJR complex structure.

### Validation on GLP-1R

GLP-1R is a class B GPCR that regulates glucose homeostasis and appetite, making it a key therapeutic target for diabetes and obesity ^44–46^. While numerous agonist peptides, including exenatide, liraglutide, semaglutide, and tirzepatide, have been successfully developed through established approaches, the development of antagonist peptides remains relatively underexplored, despite their potential for treating post-bariatric hypoglycemia (PBH) ^47^. Avexitide (exendin 9-39) ^48^ is one of the few clinically tested antagonists, highlighting both the therapeutic need and the design challenges in this area. This gap provides an opportunity to evaluate our state-specific computational framework for designing more challenging antagonist peptides.

The design pipeline follows a similar approach to that used for APJR. Peptide backbones are first identified using FoldSeek ^40^, followed by the design of antagonist peptide candidates targeting GLP-1R using an inverse folding model ^29^. The Supplementary Table **S3** shows the peptide seeds discovered via FoldSeek and filtered in the design pipeline. The initial design pool consists of approximately 5,000 GLP-1R antagonist candidates, with scores concentrated at the higher end. After applying essential filtering criteria (pLDDT > 85 and pAE < 6.5), around 1,400 peptides remained. Additionally, Filter-2 (introduced in the previous section) is applied to exclude peptides predicted by the model to be functionally inconsistent. Finally, like in the APJR experiment, corse MMGBSA scoring is used to select the top 10 peptides for experimental validation, as shown in Figure 11a. Among these 10 peptides, 4 peptides exhibit inhibitory function, with two of them demonstrating superior affinity compared to the phase-III peptide drug Avexitide (Figure 11b). The experimental affinity results are listed in Table 5.

To explore the novelty of the designed sequences, we compared the designed peptides (I3 and I4) with the sequences of the searched peptide backbones (marked with an asterisk) and Avexitide for similarity. A multiple sequence alignment was performed using MAFFT. Peptide-I3, Peptide-I4, and Avexitide function as antagonists (marked in blue), while SAR425899^49^, ZP3780^50^, and GLP-1 act as agonists (marked in red). The remaining sequences correspond to unrelated protein segments or peptides identified through the peptide backbone search method (denoted in black). Although the searched peptide backbones (marked with an asterisk) do not include known antagonists, our approach successfully generated novel antagonist peptides, Peptide-I3 and Peptide-I4. Notably, despite their higher sequence similarity to agonists such as GLP-1, both peptides exhibit antagonistic behavior. This finding highlights the accuracy of our structure-based predictions in distinguishing functional roles, even in cases of high sequence similarity.

To better understand how Peptide-I3, despite its sequence similarity to GLP-1, functions as an inhibitor, we aligned the GLP-1–GLP-1R and Peptide-I3–GLP-1R complexes and analyzed their interaction patterns (Figure 11(d-h)). As shown in Figure 11d, both GLP-1 and Peptide-I3 adopt alpha-helical conformations and occupy the same receptor binding pocket. The receptor adopts an inactive conformation, characterized by an inward displacement of the intracellular end of the TM6 domain (marked in blue) from an active state (marked in red). The differences in the interaction modes of GLP-1 and Peptide-I3 with GLP-1R can be observed in the zoomed-in views, where the common peptide residues (same amino acid type at the exact location) are labeled in black, while the differing peptide residues are marked in red and light blue, respectively. For example, in Figure 11e, the 25-th residue of GLP-1 is Trp(W) with the hydrophobic side chain. In contrast, the 25-th residue of Peptide-I3 is Asp(D) with the negatively charged side chain. As observed, the significant differences are highlighted in Figures 11e, 11g, and 11h. These differences lead to a distinct interaction pattern, as shown in Figure 11i.

**Figure 5.**
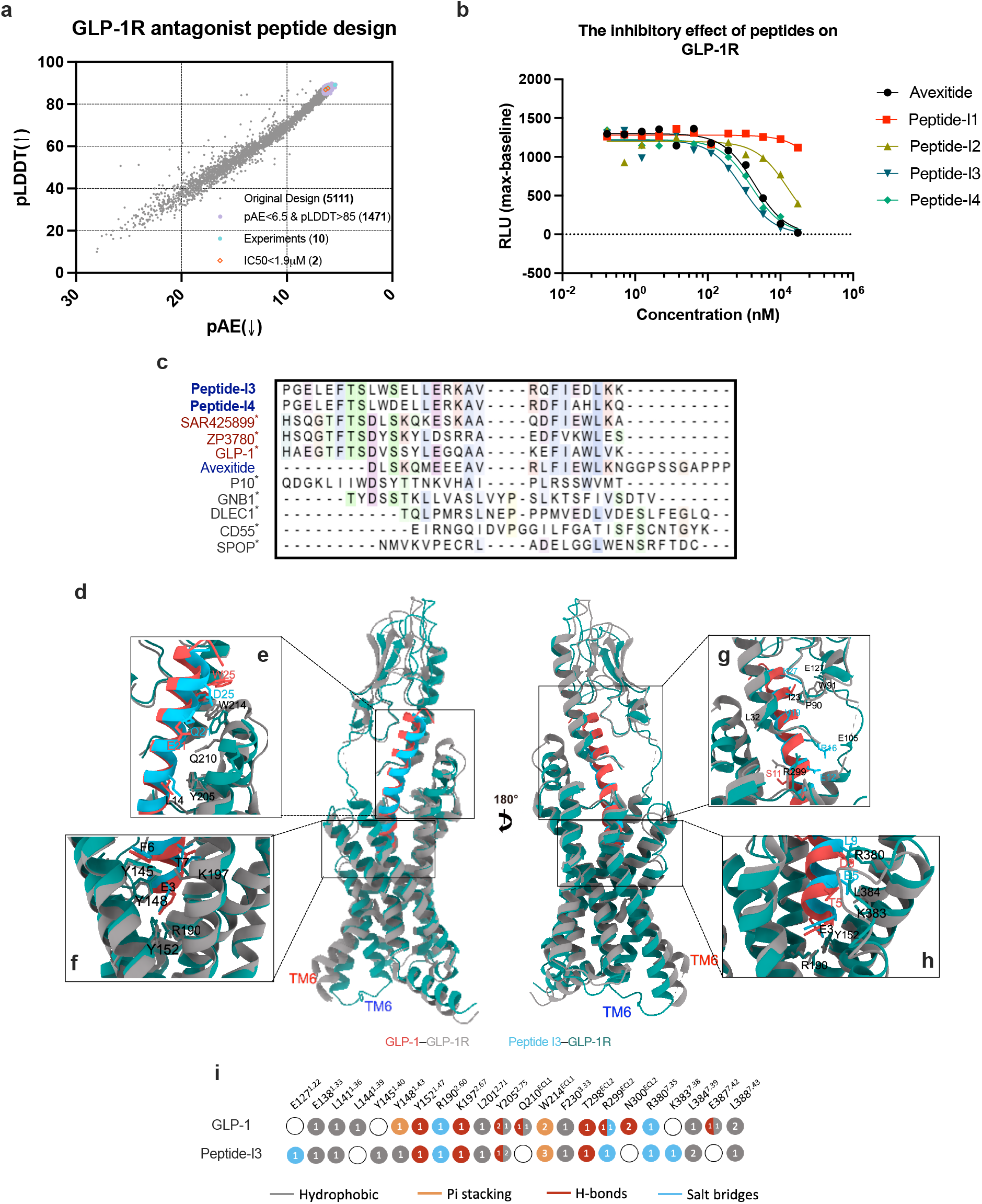
Peptides filtering and analysis for GLP-1R. **(a)** Multiple stages of computational screening. The distribution of pAE and pLDDT scores for the designed peptides is shown across various stages. **(b)** Inhibitory effects of selected peptides on GLP-1R, comparing their relative fluorescent units against varying concentrations, with Avexitide as a reference. **(c)** Sequence alignment of Peptide-I3 and Peptide-I4 with Avexitide and top seeds of the peptide backbone (marked with ^*^) used in the design pipeline. Antagonist peptides are indicated in blue, while agonist peptides are shown in red. **(d-h)** GLP-1-GLP-1R and Peptide I3-GLP-1R complexes structure alignment and zoom-in views. The GLP-1 complex is PDB **6X18**, and the Peptide-I3 complex is predicted by HF-Multistate in an inactive state. **(i)** Comparison of interaction patterns of GLP-1 agonist and Peptide-I3 with the GLP-1 receptor.

### Validation on GHSR

The growth hormone secretagogue receptor (GHSR) is a G-protein-coupled receptor that regulates the release of growth hormone and appetite ^51^. Its endogenous agonist, Ghrelin ^52^, and antagonist, LEAP2^53^, serve as key modulators of its function.

**Table 2.**
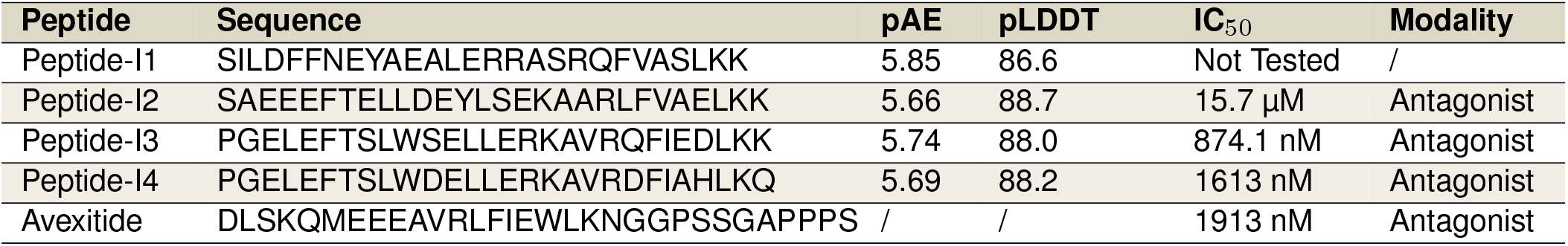
Experimental affinity and peptide modality for designed antagonist peptides targeting GLP-1R

**Figure 6.**
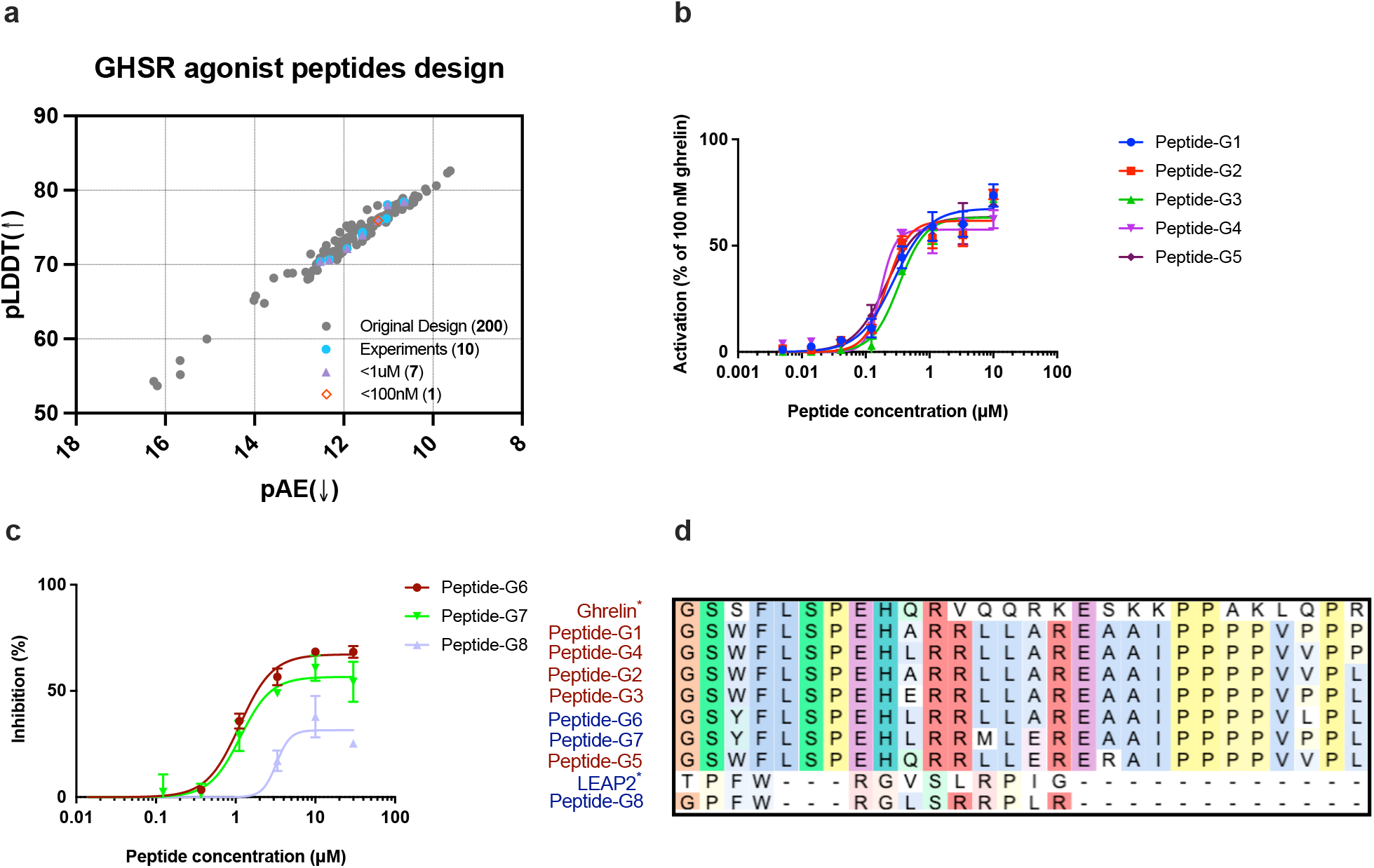
Peptides filtering and analysis for GHSR. **(a)** Multiple stages of computational screening. The distribution of pAE and pLDDT scores for the designed peptides is shown across various stages. **(b)** Dose-response curves showing the change in fluorescence intensity ratio (%) as a function of peptide concentration (µM) for various GHSR agonist peptides. **(c)** Inhibition ratios (%) of GHSR activity in response to increasing concentrations of designed GHSR antagonist peptides. **(d)** Sequence alignment of our design peptides and seeds Ghrelin and LEAP2 used in the design pipeline. Antagonist peptides are indicated in blue, while agonist peptides are shown in red.

To investigate peptide design strategies that yield well-defined binding interactions, we employed Ghrelin and LEAP2 as seed sequences. Given their established binding modes, we constrained key hotspot residues and utilized ProteinMPNN ^30^ to generate a focused library of 200 peptide candidates. Binding affinities were subsequently assessed using HF-Multistate, and candidates with a PLDDT value greater than 70 were selected. We utilize the GHSR state-transition mechanism ^52^ as an additional filtering method. The distance (dWF) WFF cluster indicates peptide activity. A shift toward the active state suggests an agonist, while a shift toward the inactive state implies antagonistic activity ^52^. The remaining candidates were further refined using coarse MMGBSA scoring like in the APJR experiment, and the top 10 peptides were selected for experimental validation (Figure 12a).

For agonist design, Figure 12b shows the majority of tested peptides exhibited EC50 values below 1 µM, with Peptide G4 demonstrating sub-100 nM potency. This suggests that structure-based design, when combined with computational scoring, can effectively identify high-affinity peptide candidates. For antagonist design, the inhibition curve in Figure 12c demonstrates that Peptide G6 to G8 exhibited an IC50 in range of 1.0-3.3 µM, effectively disrupting Ghrelin-GHSR interactions. Notably, some peptides derived from the Ghrelin scaffold displayed antagonistic activity, suggesting that HF-Multistate, when specifying GPCR state, can partially capture functional shifts in peptide design. However, unlike the APJR and GLP-1R experiments, we only utilized canonical peptide seeds Ghrelin and LEAP2 to design peptides for GHSR without FoldSeek-based diverse seeds discover process, the designed sequences are not as diverse as previous two experiments (Figure 12d). This result (Table 6) highlights the importance of diverse peptide seed backbones because current inverse folding based models may converge to several sequences given limited backbones.

**Table 3.**
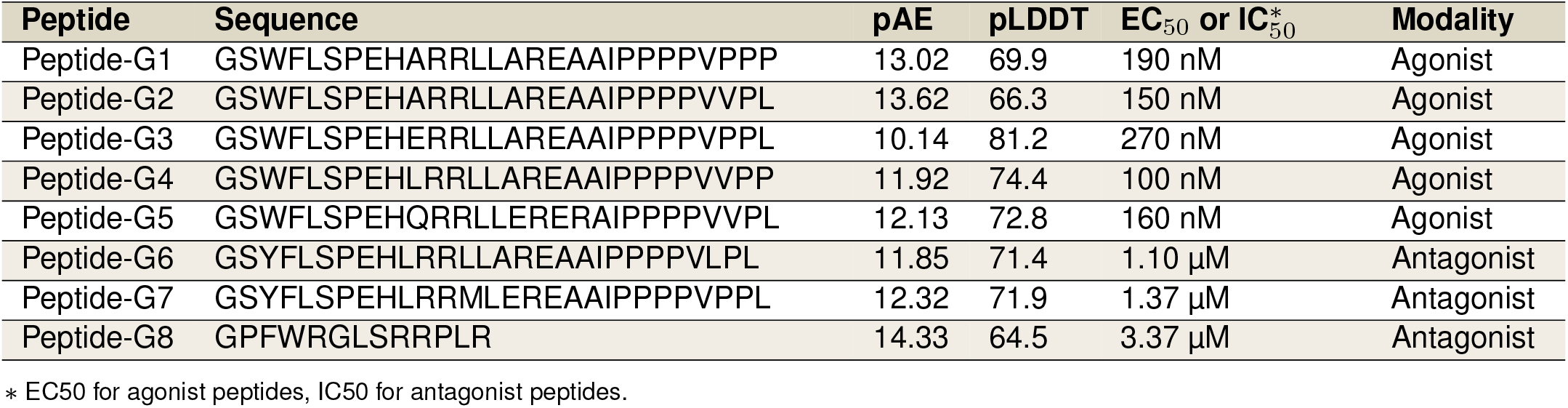
Experimental affinity and peptide modality results for designed peptides targeting GHSR

## Discussion

The development of state-specific peptides targeting G protein-coupled receptors (GPCRs) represents a significant advancement in the field of drug discovery and peptide therapeutics. GPCRs, as key mediators of cellular signaling, have long been prime targets for pharmaceutical interventions. Recent successes in GPCR-targeted peptide therapeutics underscore the potential of this approach. For instance, the FDA approval of semaglutide, a GLP-1 receptor agonist peptide for type 2 diabetes and obesity treatment, has demonstrated the clinical efficacy of peptides in modulating GPCR activity ^44;45^. Similarly, the development of dasiglucagon, a glucagon receptor agonist peptide for the treatment of severe hypoglycemia, showcases the potential of engineered peptides to improve upon natural ligands in terms of stability and efficacy ^54^. However, these successes also illuminate the challenges in peptide design, particularly in accounting for the dynamic nature of GPCRs. Traditional peptide design methods often overlook the multiple conformational states of GPCRs, which can significantly impact ligand binding and receptor activation. Our computational pipeline addresses this critical gap by explicitly incorporating the GPCR activation state into the peptide design process. By leveraging diversified seed peptide backbones, inverse folding models for sequence design, and state-specific refolding and scoring for filtering, our approach offers a more nuanced and potentially more effective method for designing GPCR-targeting peptides.

Our state-specific peptide design approach for GPCRs addresses several limitations of existing methods and offers unique advantages. Unlike conventional approaches that rely on natural peptides, phage display, or single-state computational methods, our pipeline explicitly accounts for the dynamic conformational states of GPCRs. This is a significant advancement over recent rational design methods like EvoBind ^18^, EvoPlay ^21^, or PepMLM ^23^, which do not consider GPCR state transitions. While approaches like AlphaFold-Multistate ^34^ attempt to address GPCR activation states, our experiments revealed its limitations in accurately predicting GPCR-peptide complex structures. Our pipeline overcomes this challenge by combining diversified seed peptides, inverse folding models for sequence generation, and a specifically optimized GPCR-peptide structure prediction model. This integrated approach allows us to design unconstrained, state-specific functional peptides, addressing a gap left by methods like HelixDiff ^28^, which focuses solely on structure-stable helix peptides. However, we acknowledge that the performance difference between agonist and antagonist design highlights an area for future refinement, particularly in designing peptides that must prevent receptor activation.

The success of our state-specific peptide design pipeline for GPCRs opens up exciting avenues for future research and development in the field of peptide therapeutics. While our current work has focused primarily on optimizing binding affinity, future iterations of this approach should aim to simultaneously optimize multiple pharmacological properties. Specifically, enhancing potency and extending half-lives are crucial next steps in developing peptide therapeutics with improved clinical efficacy. This could involve incorporating additional scoring functions or filters that account for factors affecting in vivo stability and receptor activation efficiency. Moreover, the versatility of our approach presents an opportunity to design peptides targeting multiple GPCR targets simultaneously, similar to the multi-target drug tirzepatide. Such multi-target peptides could offer synergistic therapeutic effects and potentially address complex disorders involving multiple GPCR pathways. Another promising direction is the integration of our pipeline with advanced peptide modification techniques, such as cyclization or the incorporation of non-natural amino acids, to further enhance the pharmacokinetic properties of the designed peptides. Furthermore, extending the application of our method to less well-characterized GPCRs could dramatically expand the scope of peptide-based drug discovery. To facilitate this, future work should focus on improving the accuracy of GPCR state prediction for receptors with limited structural data. Lastly, the incorporation of diffusion model techniques to predict all-atom structures directly leads to fewer MSA requirements ^55^, and peptides may benefit from this. By addressing these future directions, our state-specific peptide design approach has the potential to revolutionize GPCR-targeted therapeutics, leading to the development of more effective and safer peptide drugs with optimized pharmacological profiles.

## Methods

### A. Data Process

We utilized data from the Propedia v1.0 database ^332^ and GPCRdb^32^ (data released before June 30, 2021), comprising over 7,000 entries, to fine-tune HelixFold-Multimer to enhance its capabilities in predicting GPCR-peptide complex structures and improving peptide screening efficiency. The GPCRdb data distribution and quality analysis are conducted in Supplementary Section **A**. To ensure data quality and avoid redundancy, duplicate complexes shared between the two datasets were removed from Propedia during preprocessing, retaining each complex only once.

Our data processing methodology aligns with the workflow used in AlphaFold-Multimer ^20^. For both general protein-peptide and GPCR-peptide complexes, we employed the corresponding template processing pipelines from AlphaFold-Multimer ^20^ and AlphaFold-Multistate ^34^. Specifically, for GPCR templates, we selected those that matched both the same class and activation state as the target receptor. This approach ensures that the templates provide a biologically relevant and consistent structural basis for accurate predictions and evaluations.

Data from GPCRdb released after June 30, 2021^32^, filtered for 45 high-quality complexes (30 from class A and 15 from class B), was used to benchmark GPCR-peptide structure prediction across various folding models. It is essential to note that “high-quality complexes” refers to GPCR structures that are intact, enabling evaluation tools like DockQ to accurately align and calculate DockQ scores. The screening evaluation dataset, CXCR4-peptide, was initially collected in PPI-Affinity ^37^. We compiled the data list and utilized the same multistate data pipeline as mentioned in AlphaFold-Multistate ^34^ to process state-specific input features, evaluating and comparing AF-Multistate and our proposed HF-Multistate.

### B. Peptide Design Module

To generate backbone seeds, we use the FoldSeek tool ^40^ with default settings to search the Protein Data Bank for crystallized structures similar to the target structure. The search results are then parsed to identify hits that exhibit high contact density in the desired interface area, such as the groove of the GPCR protein’s extracellular region. These hits are ranked based on the number of contacts within a continuous segment, and the highest-ranking hit is selected to extract the backbone of the interface segment, which serves as the seed for protein sequence design.

For sequence design, we employ the protein inverse folding model ^29^, a computational approach that predicts amino acid sequences based on a given three-dimensional structure. In this study, we apply this model to design a diverse set of candidate peptide sequences derived from co-crystal structures of GPCR targets and peptide backbones. These peptide candidates are then evaluated using HF-Multistate to identify the most promising sequences for further investigation.

### C. State-Specific Folding Model to Select Promising Peptides

The fine-tuned HelixFold-Multistate model is employed to identify promising peptide candidates by accurately predicting their binding conformations to GPCR targets. All candidate peptides, generated by the peptide design module, are paired with state-specific GPCR targets and undergo structural prediction using this model. The resulting GPCR-peptide complex structures are rigorously assessed to ensure they align with the specified GPCR activation state. Peptides whose predicted structures do not match the target state are systematically filtered out, ensuring that only those with the highest functional relevance are retained for further analysis. In addition, building on the effectiveness of confidence metrics demonstrated by previous studies ^18;23;56–60^, we incorporate pAE interaction and peptide pLDDT metrics, derived from the folding model, to further refine the selection. A smaller pAE interaction value and a larger peptide pLDDT indicate a better peptide, contributing to a more accurate identification of high-performing candidates. This comprehensive approach increases the reliability of peptide selection for subsequent experimental validation.

### D. Experimental Validation

#### Cell lines and eagents

CHO-K1/AGTRL1/Gα15 (#M0025), CHO-K1/GHSR (#M00189) and CHO-K1/GLP1R/Gα15 (#M00451) Stable Cell Lines were purchased from GenScript (NanJing). All peptides including Apelin-13, Ghrelin and Avextide were synthesized by GenScript (NanJing). Calcium5 Assay Kit (#R8186) was purchased from Molecular Device. Probenecid (#P8761) and other chemical reagents were purchased from Sigma (Shanghai).

#### Calcium mobilization assay

CHO cells stably expressing either the human GHSR receptor or the human hAGTR1 receptor plus the Gα15 protein were digested with 0.25% trypsin and resuspended in cell culture medium and 10,000 cells of CHO-K1/AGTRL1/Gα15, 8,000 cells of CHO-K1/GHSR and 8,000 cells of CHO-K1/GLP1R/Gα15 were transferred respectively to each well of a black Costar 384-well optical bottom plate (Thermo #142761). Each plate was incubated at 37°C, 5% CO2 for 24 h. The culture media was removed from the plates, cells were loaded with Calcium 5 dye in an HBSS-based buffer containing 2.5mM probenecid in a total volume of 50 µL. Cells were incubated at 25°C for 2h avoiding light, and then stimulated with peptides, Apelin-13 or Ghrelin at various concentrations using a FLIPR Tetra plate-reader (Molecular Device). Agonist-mediated change in fluorescence (ex 488 nm, em 525 nm) was monitored in each well at 1 sec intervals for 160 sec. For antagonist assay, peptides were added at first and the fluorescence was monitored 1 sec intervals for 160 sec, and then Apelin-13, Ghrelin or GLP-1 were added and monitored for another 160 sec. Data were collected using Softmax version 4.8 (MDS Analytical Technologies) and analyzed using Prism software. Nonlinear regression analysis was performed to fit data and obtain maximum response (Emax), EC50, IC50, correlation coefficient (r2), and other parameters. All experiments were performed in duplicate at least 3 times to ensure reproducibility and data are reported as mean ± SEM.

## Data and Software Availability

HelixFold-Multistate fine-tuning data: Propedia v1.0 (http://bioinfo.dcc.ufmg.br/propedia/public/download/complex.csv), GPCRdb (https://gpcrdb.org/) (Date cutoff: June 30, 2021).

HelixFold-Multisate folding evaluation dataset: we extracted recent peptide-GPCR PDB data from GPCRdb (see Supplementary Section **B**), and collected their regions of TM domains from PDBTM (https://pdbtm.unitmp.org/) database for further state-specific analysis.

HelixFold-Multistate score screening evaluation dataset: CXCR4-peptide ^37^ (https://pubs.acs.org/doi/suppl/10.1021/acs.jproteome.2c00020/suppl_file/pr2c00020_si_001.pdf Table SI-6.1).

FoldSeek (https://github.com/steineggerlab/foldseek), ESM-IF (https://github.com/facebookresearch/esm), AlphaFold-Multistate (https://github.com/huhlim/alphafold-multistate).

## Author Contributions

Yang Xue collected and processed the data, as well as fine-tuned and evaluated the HelixFold-Multistate model. Jun Li and Yang Xue implemented the peptide design pipeline and conducted state-specific peptide designs for APJR, GHSR, and GLP-1R. Zhiyuan Chen, Lihang Liu, and Kunrui Zhu supported the base folding model, HelixFold-Multimer. Hong Wang, Jianguo Hu, and Zhi Zheng conducted the wet-lab experiments on three GPCR proteins. Jingzhou He, Huanzhang Gong, Xiangqun Li, Xiaonan Zhang, and Xiaomin Fang led the collaborative project, providing resource support and technical guidance.

## Acknowledgments

This work was supported by funding from Baidu Inc., which provided resources for AI models and computational resources. Additionally, Zonsen PepLib Biotech Inc. funded the wet lab experiments, including peptide synthesis, affinity assays, and related activities. We gratefully acknowledge their financial support.

## Footnotes

1 https://www.ebi.ac.uk/jdispatcher/msa/mafft

2 http://bioinfo.dcc.ufmg.br/propedia

