## Supplemental for "State-specific Peptide Design Targeting G Protein-coupled Receptors"

### Supplementary Information

#### A. GPCR-Peptide Structure Database

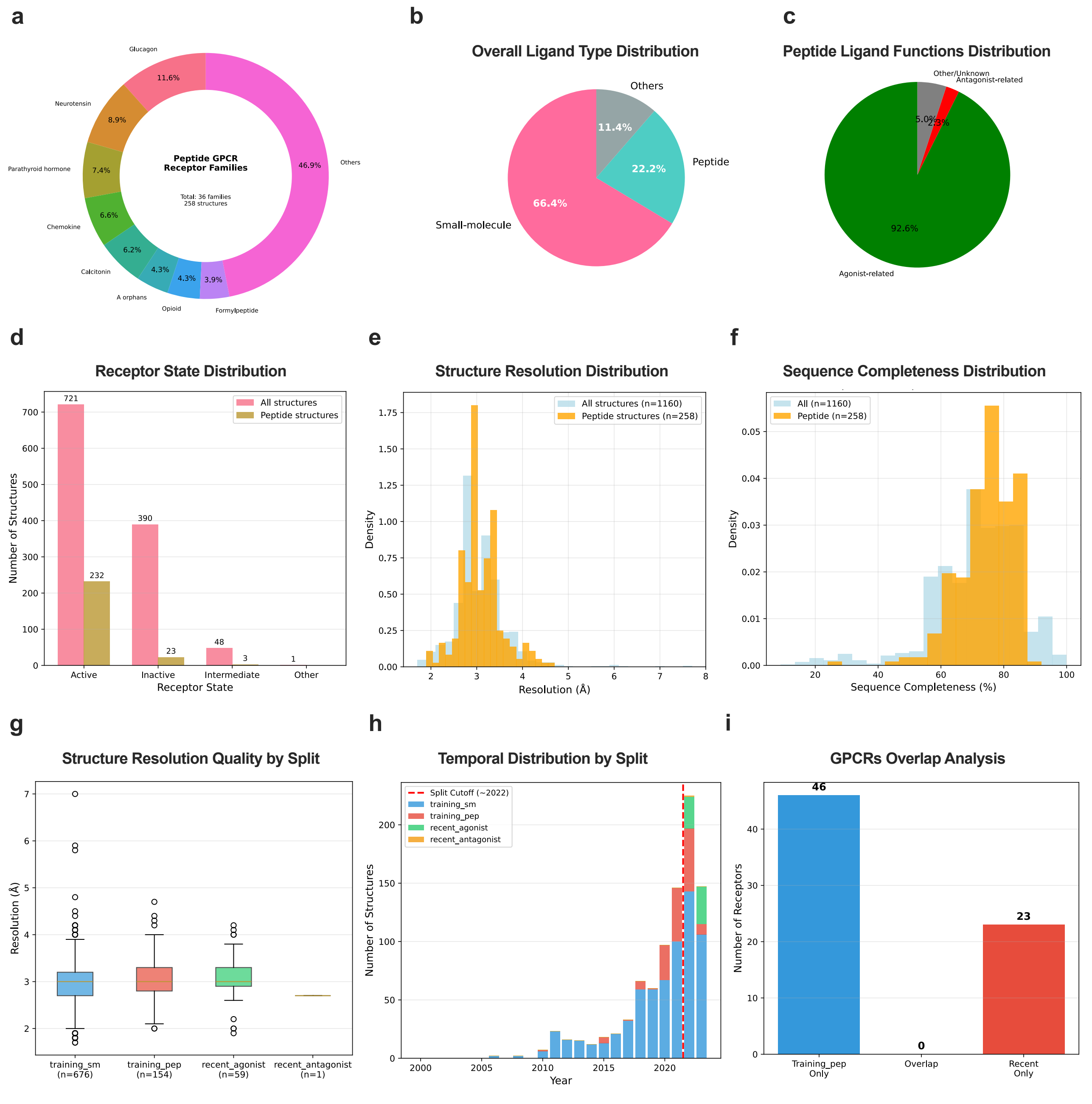


**Figure S1. GPCRdb distribution, quality and data split analysis.**

The GPCRdb Distribution, Quality and Data Split Analysis (Figure S1) provides a comprehensive overview of the GPCR-peptide database, highlighting key distributions, quality metrics and training-test data split standard. The receptor family distribution (Figure S1a) reveals the Glucagon GPCR family as the most prevalent with 11.6%, followed by the Neurotensin family at 8.9%, among a total of 36 families. Ligand type distribution (Figure S1b) indicates a strong prevalence of small-molecule ligands (66.4%), with peptides at 22.2% and others at 11.4%. Peptide ligand functions (Figure S1c) are predominantly agonist-related (92.6%), with minor antagonist (5.02%) and other/unknown roles. Receptor state distribution (Figure S1d) reveals a majority in the active state (721 structures), with fewer in inactive (390) and intermediate (23) states. Structure resolution (Figure S1e) peaks around 2-3 Å for both all (n=1160) and peptide structures (n=258), indicating good quality, while sequence completeness (Figure S1f) shows most structures (n=1160) are highly complete (>80%), with peptide structures (n=258) following a similar trend.

We divided the GPCRdb dataset into four groups: training_sm (GPCR-small molecule), training_pep (GPCR-peptide), recent agonist, and recent antagonist. For the training_sm set, we utilize only GPCR structures for training. Recent agonist and antagonist consist of GPCR-peptide PDBs allocated as an evaluation set. We filtered the evaluation set by excluding PDBs from the most recent years (Figure S1h) and ensuring no GPCR type overlap with the training_pep set (Figure S1i).

The initial evaluation set of 60 PDBs was filtered for quality. Fifteen PDBs failed DockQ analysis due to sequence completeness issues and were removed. The remaining 45 high-quality, peptide-bound GPCR structures were grouped by GPCR class as listed below:

**Class A:** 7SK7, 7VFX, 7SK4, 7SK5, 7F6G, 7W0P, 7Y67, 7XA3, 7YJ4, 7SK3, 7SK6, 7YKD, 7XWO, 7SK8, 7Y66, 7XNA, 7YON, 7Y1F, 7Y64, 8HK5, 8HK2, 8F7Q, 8H0Q, 8F7W, 8DWC, 8F7X, 8DWG, 8H0P, 8F7R, 8IA8.

**Class B:**. 7VVN, 7VVL, 7VVK, 7VVJ, 7VVM, 8FLU, 8F0J, 8F0K, 8FLR, 8HA0, 8F2B, 8HAF, 8HAO, 8F2A, 8FLQ.

Of these 45 structures, only one PDB (7XNA) is in an antagonist-bound, inactive state. Due to this scarcity, a meaningful comparison of GPCR-peptide structure prediction performance between different functional states was not possible and has been omitted from our report.

#### B. Additional Results for GPCR-peptide Structure Prediction

To evaluate the GPCR-peptide interaction, Figure S2 presents a box and whisker plot illustrating DockQ and iRMS scores for recent PDBs of GPCR class A (Figure S2 a,b) and GPCR class B (Figure S2 c,d) as supplementary information to Figure 2. This visualization clearly highlights that some failure cases result in extended whiskers. Overall, the interaction performance of HF-Multistate, in terms of DockQ and iRMS, is comparable to the best-performing model, AF-Multimer. In contrast, AF-Multistate exhibits a notable decline in performance.

To evaluate the GPCR activation state, Figure S3 and Figure S4 compare HF-Multistate with AF-Multimer, AF-Multistate, and HF-Multimer in terms of TM RMSDs. Additionally, we define **success rate** as the proportion of cases where the TM RMSD is less than 2Å and **beat rate** as the percentage of cases where the TM RMSD is lower than that of the compared model. As shown, HF-Multistate consistently outperforms the other models.

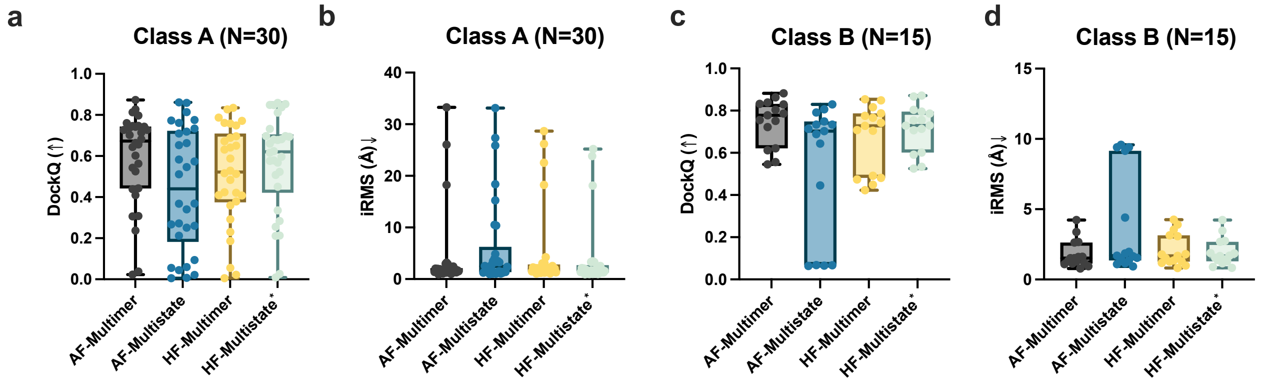


**Figure S2. Comparing activation state prediction performance for GPCR class A and B using DockQ and iRMS metrics.** **(a)** and **(b)** correspond to GPCR Class A, while **(c)** and **(d)** pertain to GPCR Class B.

#### C. FoldSeek discovered seed peptides from APJR and GLP-1R

To identify seed peptides for APJR, we begin by gathering APJR-related PDBs and classifying them into active (7WOL, 7WOM, 7WON, 7WOP) and inactive (5VBL, 6KNM, 7SUS) states. In the initial phase, FoldSeek is employed with default settings to detect peptide seeds, producing 781, 836, and 816 seeds from inactive states, and 830, 713,

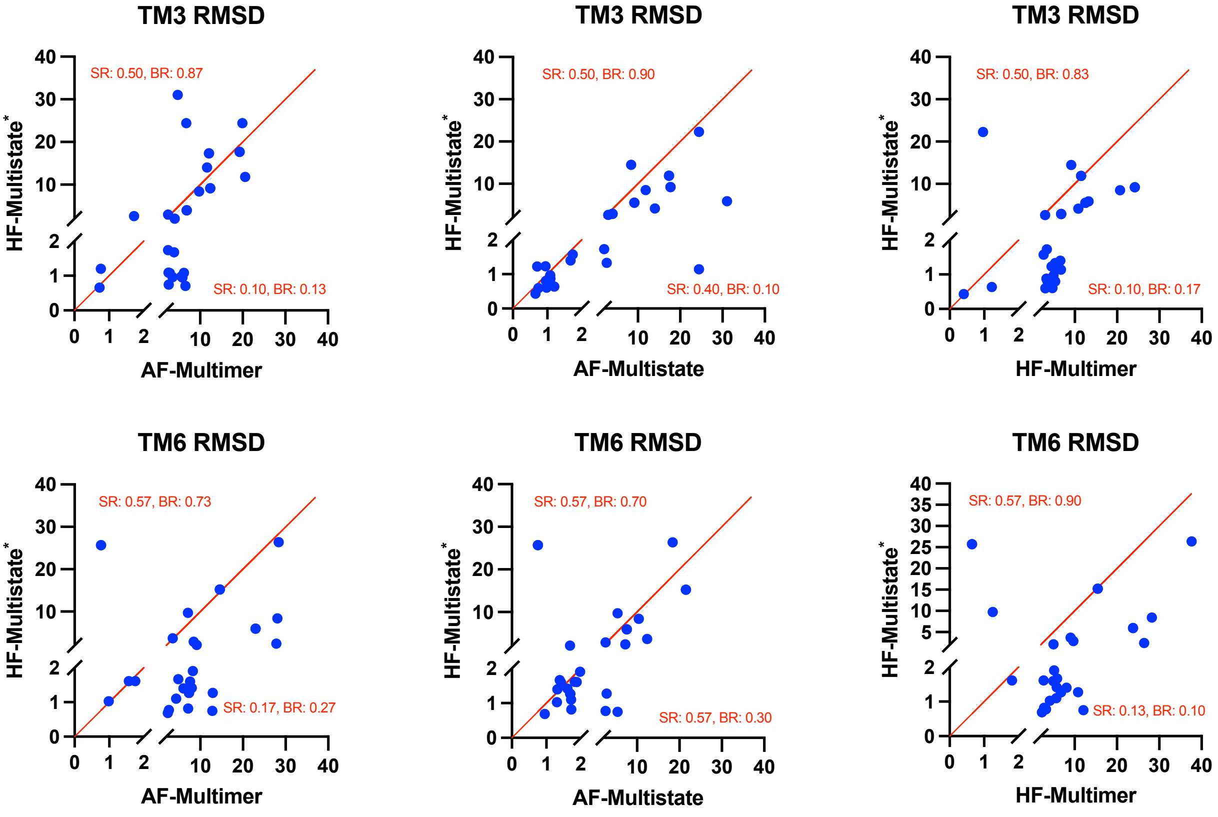


**Figure S3. Comparing activation state prediction performance for GPCR class A on TM3 and TM6 domains.** For GPCR class A proteins, their activation states are mainly determined by TM3 and TM6 domains. The corresponding RMSD metrics reflect that our proposed HF-Multistate model outperforms baselines in terms of SR (success rate) and BR (beat rate).

786, 705, and 686 seeds from active states, respectively. The subsequent phase involves consolidating these seeds and removing duplicates to enhance the dataset's quality. The first refinement applies criteria, including a sequence length of 3-25 and a contact density (average number of contacts per peptide residue) > 2, reducing the set to 72 active and 48 inactive seeds. Finally, the second refinement visualizes and confirms binding to EOD, narrowing the selection to 21 active (Table S1) and 16 inactive (Table S2) peptide seeds, thereby concluding the discovery process.

For GLP-1R, we concentrate exclusively on the inactive state (6LN2, 6KK1, 6KJV, 6KK7, 5VEW, 5VEX), employing a comparable methodology to that used for APJR, except with a maximum seed length of 30 for GLP-1R. This process yields a selection of 13 inactive peptide seeds (Table S3).

#### D. Peptide-A6 and APJR Groove I region interaction stability analysis

We used gmx_MMPBSA[1] to analyze the energy decomposition for residues in the Groove I region interacting with peptide ligand residues. The total interaction energy and its component breakdown are illustrated in Figure S5. The low interaction energy between all pairs indicates a stable interaction between Peptide-G6 and APJR at the Groove I region.

[1] https://github.com/Valdes-Tresanco-MS/gmx_MMPBSA

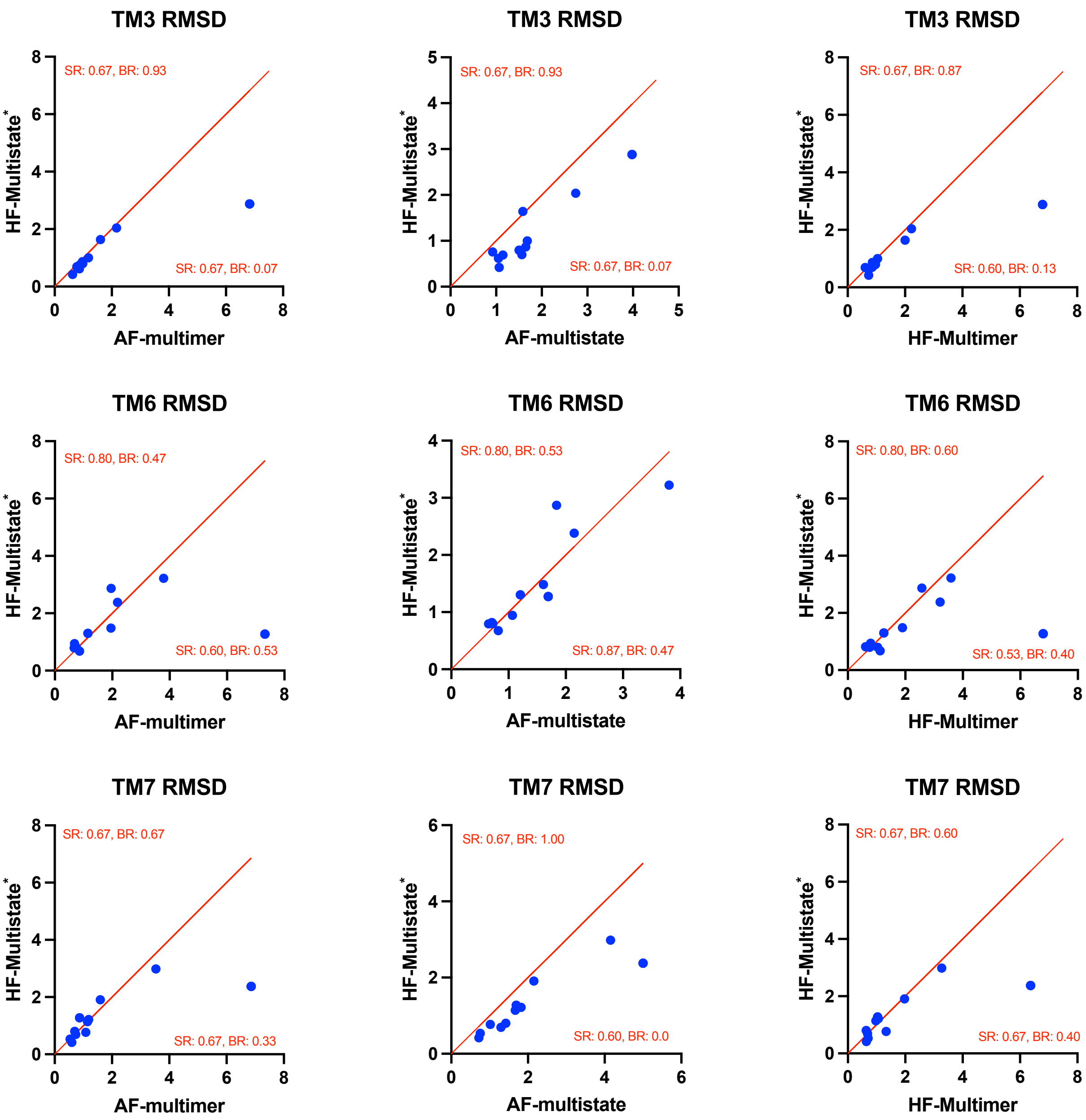


**Figure S4. Comparing activation state prediction performance for GPCR class B on TM3, TM6 and TM7 domains.** For GPCR class B proteins, their activation states are mainly determined by TM3, TM6 and TM7 domains. Unlike GPCR class A (Figure S3), there is less PDB data for class B, and the margin between HF-Multistate and baseline models is smaller.

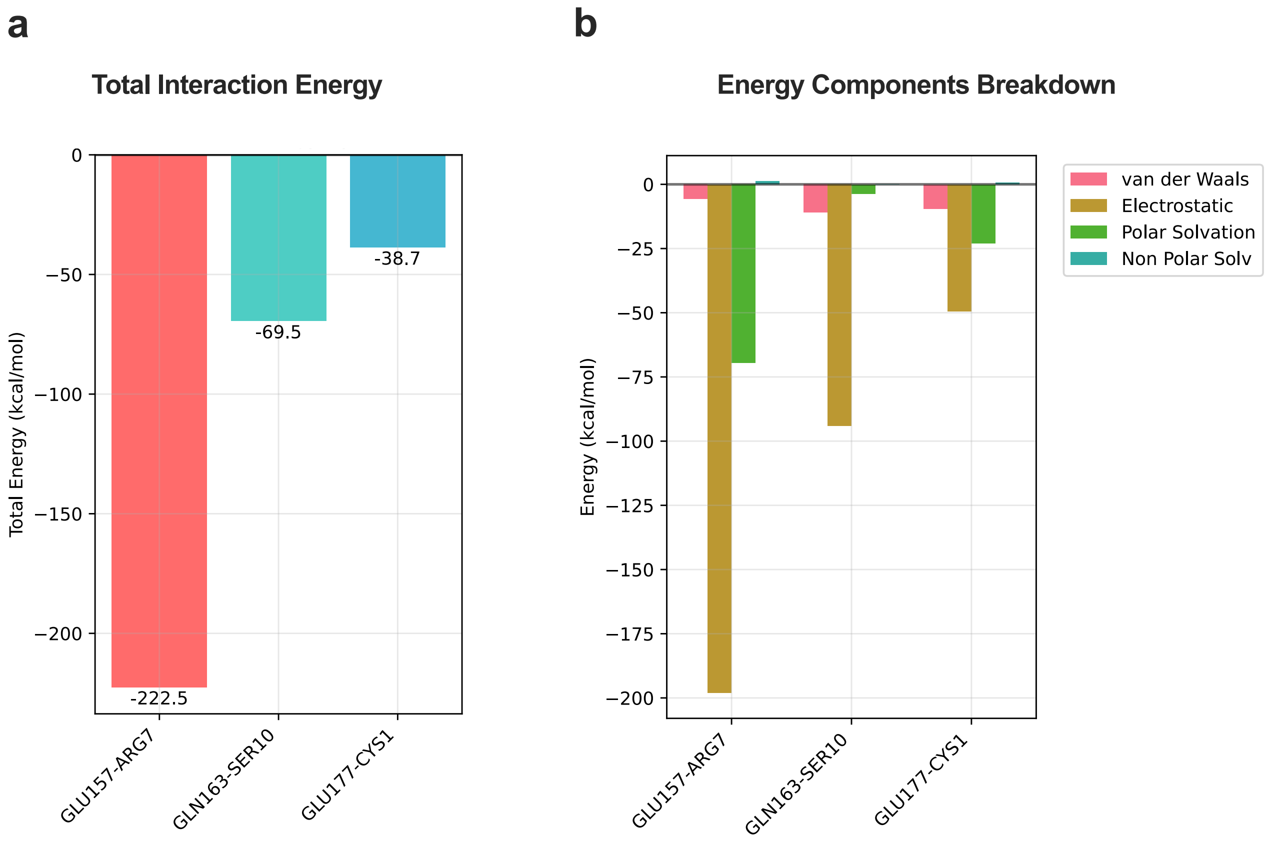


**Figure S5. Total interaction energy and components breakdown for Peptide-G6 and APJR interaction residue pairs at Groove I region.**

| PDB | target chain | target identity | seed chain | seed range | seed sequence |
|---|---|---|---|---|---|
| 4XT1 | A | 0.298 | B | 2-19 | HHGVTKCAITCSKMTSKI |
| 6JOD | A | 0.294 | B | 1-8 | DRVYIHPF |
| 6KNM | B | 0.834 | A | 99-119 | PRAGIESGAYCKWNMKDSGSW |
| 6LFO | R | 0.323 | D | 2-22 | AKELRCQCIKTYSKPFHPKFI |
| 6WWZ | R | 0.258 | C | 1-20 | ASNFDCCLGYTDRILHPKFI |
| 7O7F | C | 0.276 | I | 2-26 | PPGDIVLACCFAYIARPLPRAHIKE |
| 7SK3 | A | 0.272 | B | 1-25 | KPVSLSYRCPCRFFESHVARANVKH |
| 7SK6 | A | 0.277 | B | 0-16 | LRHQSLSYRCPCRFFES |
| 7T10 | R | 0.293 | P | 1-14 | AGCKNFFWKTFTSC |
| 7VLA | R | 0.303 | L | 27-51 | HFAADCCTSYISQSIPCSLMKSYFE |
| 7W0O | R | 1.000 | D | 18-32 | LQRRCMPLHSRVPFP |
| 7WIC | R | 0.282 | L | 1-14 | AGCKNFFWKTFTSC |
| 7WQ3 | R | 0.257 | L | 2-14 | WTLNSAGYLLGPH |
| 7WQ4 | R | 0.295 | L | 2-12 | WTLNSAGYLLG |
| 7X9B | R | 0.245 | P | 21-35 | YSALRHYINLITRQR |
| 7XA3 | R | 0.271 | L | 1-18 | QPDAINAPVTCCYNFTNR |
| 7XBD | A | 0.280 | F | 2-14 | WTLNSAGYLLGPH |
| 7XJJ | E | 0.247 | C | 2-16 | WTLNSAGYLLGPHAV |
| 7XJL | F | 0.275 | A | 1-14 | NWTPQAMLYLKGAQ |
| 7Y1F | R | 0.289 | F | 1-9 | YGGFLRRIR |
| 8F7W | R | 0.269 | P | 1-8 | YGGFLRRI |

**Table S1.** Peptide seeds discovered by FoldSeek and filtered for APJR agonist peptide design

| PDB | target chain | target identity | seed chain | seed range | seed sequence |
|---|---|---|---|---|---|
| 4XT1 | A | 0.246 | B | 2-19 | HHGVTKCAITCSKMTSKI |
| 4XT3 | A | 0.250 | B | 2-18 | HHGVTKCNITCSKMTSK |
| 6KNM | B | 1.000 | A | 99-119 | PRAGIESGAYCKWNMKDSGSW |
| 6LFO | R | 0.286 | D | 2-22 | AKELRCQCIKTYSKPFHPKFI |
| 6WWZ | R | 0.232 | C | 1-20 | ASNFDCCLGYTDRILHPKFI |
| 7F6I | A | 0.254 | L | 1-10 | KRPPGFSPFR |
| 7O7F | C | 0.230 | I | 2-26 | PPGDIVLACCFAYIARPLPRAHIKE |
| 7SK3 | A | 0.248 | B | 1-25 | KPVSLSYRCPCRFFESHVARANVKH |
| 7SK6 | A | 0.277 | B | 0-16 | LRHQSLSYRCPCRFFES |
| 7VLA | R | 0.245 | L | 27-51 | HFAADCCTSYISQSIPCSLMKSYFE |
| 7WIC | R | 0.241 | L | 1-14 | AGCKNFFWKTFTSC |
| 7WQ3 | R | 0.229 | L | 2-14 | WTLNSAGYLLGPH |
| 7XA3 | R | 0.231 | L | 1-18 | QPDAINAPVTCCYNFTNR |
| 7XJL | F | 0.226 | A | 1-14 | NWTPQAMLYLKGAQ |
| 8F7W | R | 0.250 | P | 1-8 | YGGFLRRI |
| 8HK2 | A | 0.257 | D | 62-77 | ELRRQHARASHLGLAR |

**Table S2.** Peptide seeds discovered by FoldSeek and filtered for APJR antagonist peptide design

| PDB | target chain | target identity | seed chain | seed range | seed sequence |
|---|---|---|---|---|---|
| 4HJ0 | B | 0.416 | C | 100-108 | PIVGAPTDY |
| 5XEZ | A | 0.676 | C | 316-327 | HYDILTGYNYYY |
| 5YQZ | R | 0.674 | P | 1-28 | SQGTFTSEYSKYLDSRRAQDFVKWLLNT |
| 7DUQ | R | 0.839 | P | 1-30 | HAEGTFTSDVSSYLEGQAAKEFIAWLVKGR |
| 7KI1 | R | 0.842 | P | 1-30 | HAEGTFTSDVSSYLEGQAAKEFIAWLVKAR |
| 7LLY | R | 0.786 | P | 1-26 | HSQGTFTSDYSKYLDSRRAQDFVQWL |
| 7RGP | R | 0.830 | P | 1-30 | YAEGTFTSDYSIALDKIAQKAFVQWLIAGG |
| 7RKN | R | 0.118 | L | 1-30 | QHHGVTKCNITCSKMTSKIPVALLIHYQQN |
| 7RTB | R | 0.573 | P | 1-30 | YAEGTFTSDYSIYLDKQAAAEFVNWLLAGG |
| 7S1M | R | 0.836 | P | 1-29 | HGEATFTSDLSKQMEEEAVRLFIEWLKNG |
| 7VAB | R | 0.416 | P | 1-27 | YAEGTFTSDYSIALDKIAQKAFVQWLI |
| 8JIR | R | 0.842 | P | 1-29 | HSQGTFTSDLSKQKESKAAQDFIEWLKAG |
| 8JIS | R | 0.841 | P | 1-29 | HSQGTFTSDYSKYLDEQAAKEFIAWLMNT |

**Table S3.** Peptide seeds discovered by FoldSeek and filtered for GLP-1R antagonist peptide design
